# Consolidation alters motor sequence-specific distributed representations

**DOI:** 10.1101/376053

**Authors:** Basile Pinsard, Arnaud Boutin, Ella Gabitov, Ovidiu Lungu, Habib Benali, Julien Doyon

## Abstract

FMRI studies investigating the acquisition of sequential motor skills in humans have revealed learning-related functional reorganizations of the cortico-striatal and cortico-cerebellar motor systems in link with the hippocampus. Yet, the functional significance of these activity level changes is not fully understood as they convey the evolution of both sequence-specific knowledge and unspecific task expertise. Moreover, these changes do not specifically assess the occurrence of learning-related plasticity. To address these issues, we investigated local circuits tuning to sequence-specific information using multivariate distances between patterns evoked by consolidated or newly acquired motor sequences production. Results reveal that representations in dorsolateral striatum, prefrontal and secondary motor cortices are greater when executing consolidated sequences than untrained ones. By contrast, sequence representations in the hippocampus and dorsomedial striatum are less engaged. Our findings show, for the first time in humans, that complementary sequence-specific motor representations evolve distinctively during critical phases of skill acquisition and consolidation.

## Introduction

Animals and humans are able to acquire and automatize new sequences of movements, hence allowing them to expand and update their repertoire of complex goal-oriented motor actions for long-term use. To investigate the mechanisms underlying this type of procedural memory in humans, a large body of behavioral studies has used motor sequence learning (MSL) tasks designed to test the ability to perform temporally ordered and coordinated movements, learned either implicitly or explicitly and has assessed their performances in different phases of the acquisition process (Korman et al. 2003; Abrahamse et al. 2013;Diedrichsen and Kornysheva 2015;Verwey et al. 2015). While practice of an explicit MSL task leads to substantial within-session execution improvements, there is now ample evidence indicating that between-session maintenance, and even increases, in performance can be observed after a night of sleep (Nettersheim et al. 2015;Landry et al. 2016), while performance are unstable and tends to decay during an equal period of wake (Doyon et al. 2009b;Brawn et al. 2010;Nettersheim et al. 2015;Landry et al. 2016). Therefore, it is thought that sleep favors reprocessing of the motor memory trace, thus promoting its consolidation for long-term skill proficiency (Fischer et al. 2002;see King et al. 2017;Doyon et al. 2018 for recent in-depth reviews).

Functional magnetic resonance imaging (fMRI) studies using General-Linear-Model (GLM) contrasts of activation have also revealed that MSL is associated with the recruitment of an extended network of cerebral (Hardwick et al. 2013), cerebellar and spinal regions (Vahdat et al. 2015), whose contributions differentiate as learning progresses (Karni et al. 1998; Dayan and Cohen 2011;Doyon et al. 2018). In fact, critical plastic changes (Ungerleider et al. 2002; Doyon and Benali 2005) are known to occur within the initial training session, as well as during the offline consolidation phase, the latter being characterized by a functional “reorganization” of the nervous system structures supporting this type of procedural memory function (Rasch and Born 2008;Born and Wilhelm 2012;Albouy et al. 2013b;Bassett et al. 2015;Dudai et al. 2015;Fogel et al. 2017;Vahdat et al. 2017). More specifically, MSL practice is known to activate a cortical, associative striatal and cerebellar motor network which is assisted by the hippocampus during the initial “fast-learning” phase (Albouy et al. 2013b). Yet, when approaching asymptotic behavioral performance after longer practice, activity within the hippocampus and cerebellum decreases while activity within the sensorimotor striatum increases (Doyon et al. 2002), both effects conveying the transition to the “slow-learning” phase. The same striatal regions are reactivated during sleep spindles (Fogel et al. 2017) contributing to the progressive emergence of a reorganized network (Debas et al. 2010; Vahdat et al. 2017), which is further stabilized when additional MSL practice extending over multiple days is separated by consolidation periods (Lehéricy et al. 2005).

A critical issue typically overlooked by previous MSL neuroimaging research using GLM-based activation contrasts, however, is that learning-related changes in brain activity do reflect the temporal evolution of recruited processes during blocks of practice, only some of which may be specifically related to plasticity induced by MSL. For instance, increases in activity could not only signal a greater implication of the circuits specialized in movement sequential learning *per se*, but could also result from the inherent faster execution of the motor task. Likewise, a decrease in activity could either indicate some form of optimization and greater efficiency of the circuits involved in executing the task (Wu et al. 2004), or could show the reduced recruitment of non-specific networks supporting the acquisition process. Therefore, even with the use of control conditions to dissociate sequence-specific from non-specific processes (Orban et al. 2010), the observed large-scale activation differences associated with different learning phases do not necessarily provide direct evidence of plasticity related to the processing of a motor sequence-specific representation (Berlot et al. 2018). Furthermore, it is also conceivable that these plastic changes could even occur locally without significant changes in the GLM-based regional activity level. Finally, in most studies investigating the neural substrate mediating the consolidation process of explicit MSL, the neural changes associated with this mnemonic mechanism are assessed by contrasting brain activity level of novice participants between their initial training and a delayed practice session, Therefore, they measure not only plasticity for sequence-specific (e.g. optimized chunks), but also task-related expertise (e.g. habituation to experimental apparatus, optimized execution strategies, attentional processes). The latter expertise is notably observed when participants practice two motor sequences in succession and the initial performance during sequence execution is significantly better for the subsequent than for the first sequence.

To address these specificity limitations, multivariate pattern analysis (MVPA) has been proposed to evaluate how local patterns of activity are able to reliably discriminate between stimuli or evoked memories of the same type over repeated occurrences, hence allowing to test information-based hypotheses that GLM contrasts cannot inquire (Hebart and Baker 2017). In the MSL literature, only a few studies have used such MVPA approaches to identify the regions that specialize in processing the representation of learned motor sequences (Wiestler et al. 2011;Wiestler and Diedrichsen 2013;Kornysheva and Diedrichsen 2014;Nambu et al. 2015;Yokoi et al. 2017). These studies, however, mainly focused on extensively practiced sequences over multiple training sessions across multiple days. For instance, in a recent study covering dorsal cerebral cortices only (Wiestler and Diedrichsen 2013), cross-validated classification accuracy was measured separately on activity patterns evoked by the practice of trained and untrained sets of sequences. The authors showed that the extended training increased sequence discriminability in a network spanning bilaterally the primary and secondary motor as well as parietal cortices. In another study (Nambu et al. 2015) that aimed to analyze separately the preparation and execution of sequential movements, representations of extensively trained sequences were identified in the contralateral dorsal premotor and supplementary motor cortices during preparation, while representations related to the execution were found in the parietal cortex ispilater-ally, the premotor and motor cortices bilaterally as well as the cerebellum. In both studies, the regions carrying sequence-specific representations overlapped only partly with those identified using GLM-based measures, hence illustrating the fact that coarser differences in activation between novel and trained sequences does not necessarily provide evidence of plasticity for sequential information. However, the classification-based measures they used may have biased their parametric statistical results by violating both the normality assumption and theoretical null-distribution (Allefeld et al. 2015; Combrisson and Jerbi 2015; Jamalabadi et al. 2016; Varoquaux 2017) and may have thus been suboptimal in detecting representational changes (Walther et al. 2016).

As a part of a larger research program, the present study aimed to address both the critical issues overlooked by previous research investigating the early phases of MSL consolidation with GLM-based approach described above, as well as the limitations encountered when using classifier-based MVPA methods. Specifically, we employed a recently developed MVPA approach (Nili et al. 2014) that is unbiased and more sensitive to continuous representational changes (Walther et al. 2016), such as those that occur in the early stage of MSL and consolidation (Albouy et al. 2013c). Our experimental manipulation allowed to isolate sequence-specific plasticity, by extracting patterns evoked through practice of both consolidated and new sequences at the same level of task expertise and by computing this novel multivariate distance metric using a searchlight approach over the whole brain in order to cover cortical and subcortical regions critical to MSL. Based on theoretical models (Albouy et al. 2013b; Doyon et al. 2018) derived from imaging and invasive animal studies, we hypothesized that offline consolidation following training would induce greater cortical and striatal as well as weaker hippocampal sequence-specific representations.

## Results

To investigate changes in the neural representations of motor sequences occurring during learning, young healthy participants (n=18) practiced two 5-element sequences of finger movements (executed through button presses) separately on two consecutive days. On the third day, participants were required to execute again the same two sequences, then considered to be consolidated, together with two new 5-element untrained sequences. This practice session consisted in 64 pseudo-randomly ordered short blocks split in two runs, with 16 blocks of each sequence. All four sequences were executed using their non-dominant left hand while functional MRI data was acquired.

### Behavioral performance

We analyzed the behavioral performance related to the four different sequences using a repeated-measure mixed-effects model. As expected, new sequences were performed more slowly (*β* = .365, *SE* = 0.047,*p* < .001) and less accurately (*β* = −0.304, *SE* = 0.101,*p* < .001) than the consolidated ones. Significant improvement across blocks was observed for new sequences as compared to consolidated sequences in term of change of speed (*β* = −0.018,SE = 0.002,*p* < .001), thus showing an expected learning curve visible in fig. 1. Yet accuracy did not show significant improvement (*β* = 0.014, *SE* = 0.010,*p* = 0.152) likely explained by the limited precision of this measure that ranges discretely from 0 to 5. By contrast, the consolidated sequences did not show significant changes in speed (*β* = −0.006,*SE* = 0.005,*p* = 0.192) nor accuracy (*β* = −0.006,*SE* = 0.057,*p* = 0.919), the asymptotic performances being already reached through practice and the consolidation process.

**Figure 1.**
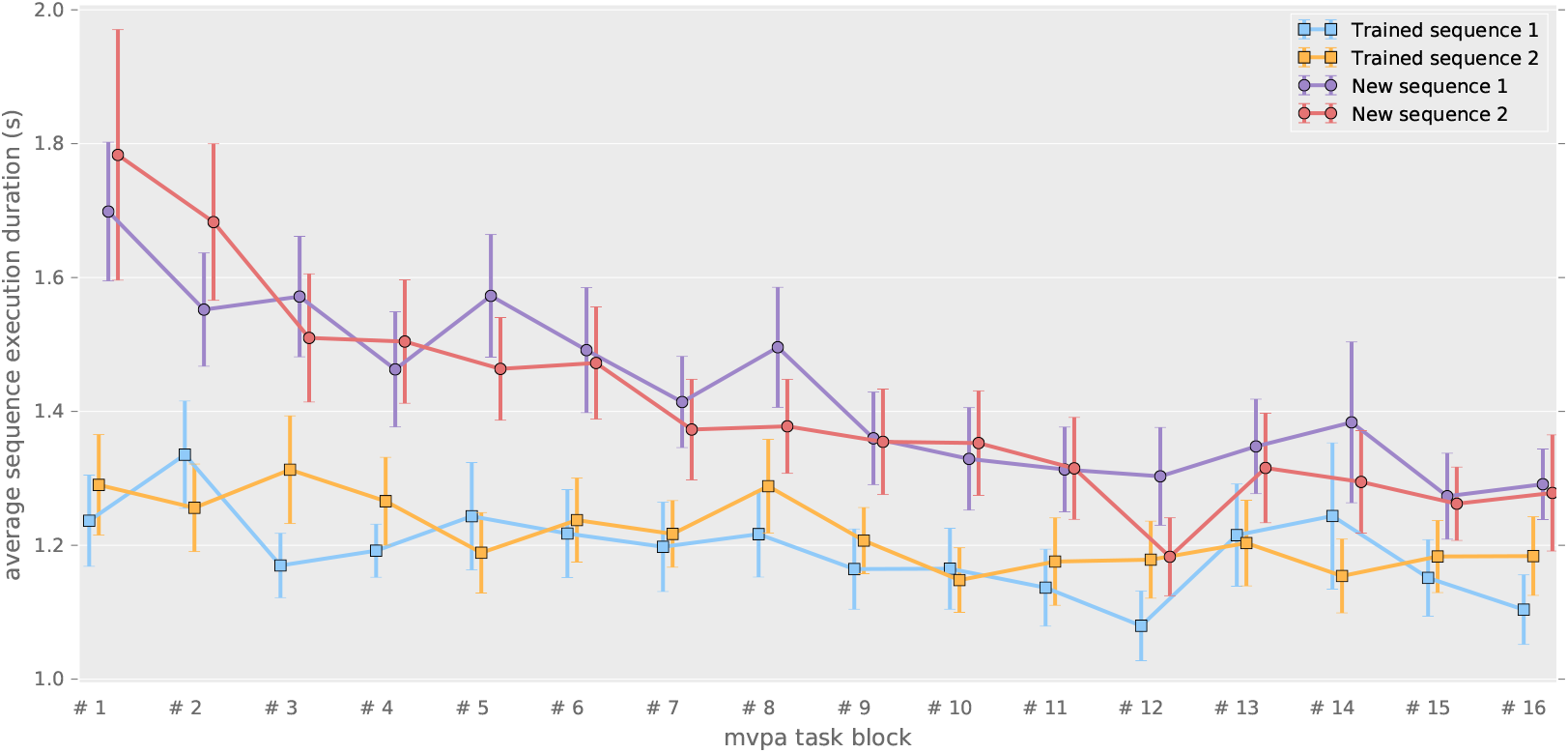
Correct sequence durations (average and standard error of the mean) across the MVPA task blocks.

Importantly, there were also no significant differences between the two consolidated sequences in term of speed (*β* = 0.031, *SE* = 0.026, *p* = 0.234) and accuracy (*β* = −0.030, *SE* = 0.111,p = 0.789), nor between the two new sequences speeds (*β* = 0.025, *SE* = 0.045,*p* = 0.577) and accuracies (*β* = −0.245, *SE* = 0.138,*p* = 0.076).

### A common distributed network for sequence representation irrespective of learning stage

From the preprocessed functional MRI data we extracted patterns of activity for each block of practice, and computed a cross-validated Mahalanobis distance (Nili et al. 2014; Walther et al. 2016) using a Searchlight approach (Kriegeskorte et al. 2006) over brain cortical surfaces and subcortical regions of interest. Such multivariate distance, when positive, demonstrate that there is a stable difference in activity patterns between the conditions compared, and thus reflect the level of discriminability between these conditions. To assess true patterns and not mere global activity differences, we computed this discriminability measure for sequences that were at the same stage of learning, thus separately for consolidated and new sequences. From the individual discriminability maps, we then measured the prevalence of discriminability at the group level, using non-parametric testing with a Threshold-Free-Cluster-Enhancement approach (TFCE) (Smith and Nichols 2009) to enable locally adaptive cluster-correction.

To extract the brain regions that show discriminative activity patterns for specific sequence during both learning stages, we then submitted these separate group results for the consolidated and new sequences to a minimum-statistic conjunction. A large distributed network (fig. 2) displayed significant discriminability, including the primary visual, as well as the posterior parietal, primary and supplementary motor, premotor and dorsolateral prefrontal cortices.(see the statistical maps for each learning stage separately in the Supplementary material (fig. S1,fig. S2).

**Figure 2.**
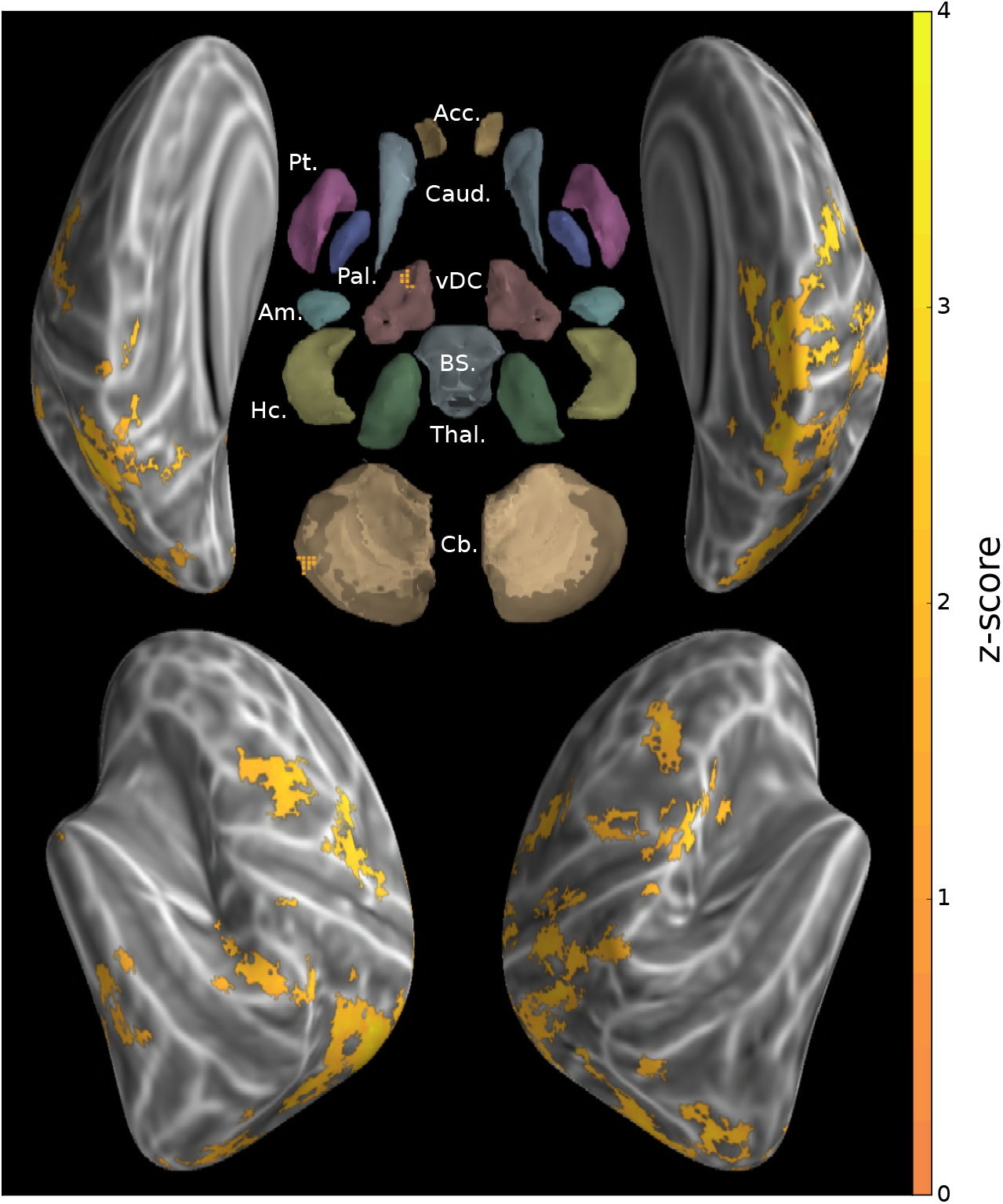
Group searchlight conjunction of new and consolidated sequences discriminability maps (z-score thresholded at *p* < .05 TFCE-cluster-corrected) showing a large distributed cortical network showing sequence disciminative patterns at both learning stages; Regions of interest with Freesurfer colors: Acc.:Accumbens; Pt.:Putamen; Caud.:Caudate; Pal.:Pallidum; vDC:ventral Diencephalon; Am.:Amygdala; Hc.:Hippocampus; Thal.:Thalamus; Cb.:Cerebellum; BS:brain-stem

### Reorganization of the distributed sequence representation after memory consolidation

In order to evaluate the reorganization of sequence representation undergone by consolidation at the group level, the consolidated and new sequence discriminability maps from all participants were submitted to a non-parametric pairwise t-test with TFCE. To ascertain that a greater discriminability in one stage versus the other was supported by a significant level of discriminability within that stage, we then calculated the conjunction of the contrast maps with the consolidated and new sequences group results, respectively with the positive and negative contrast differences (fig. 3).

**Figure 3.**
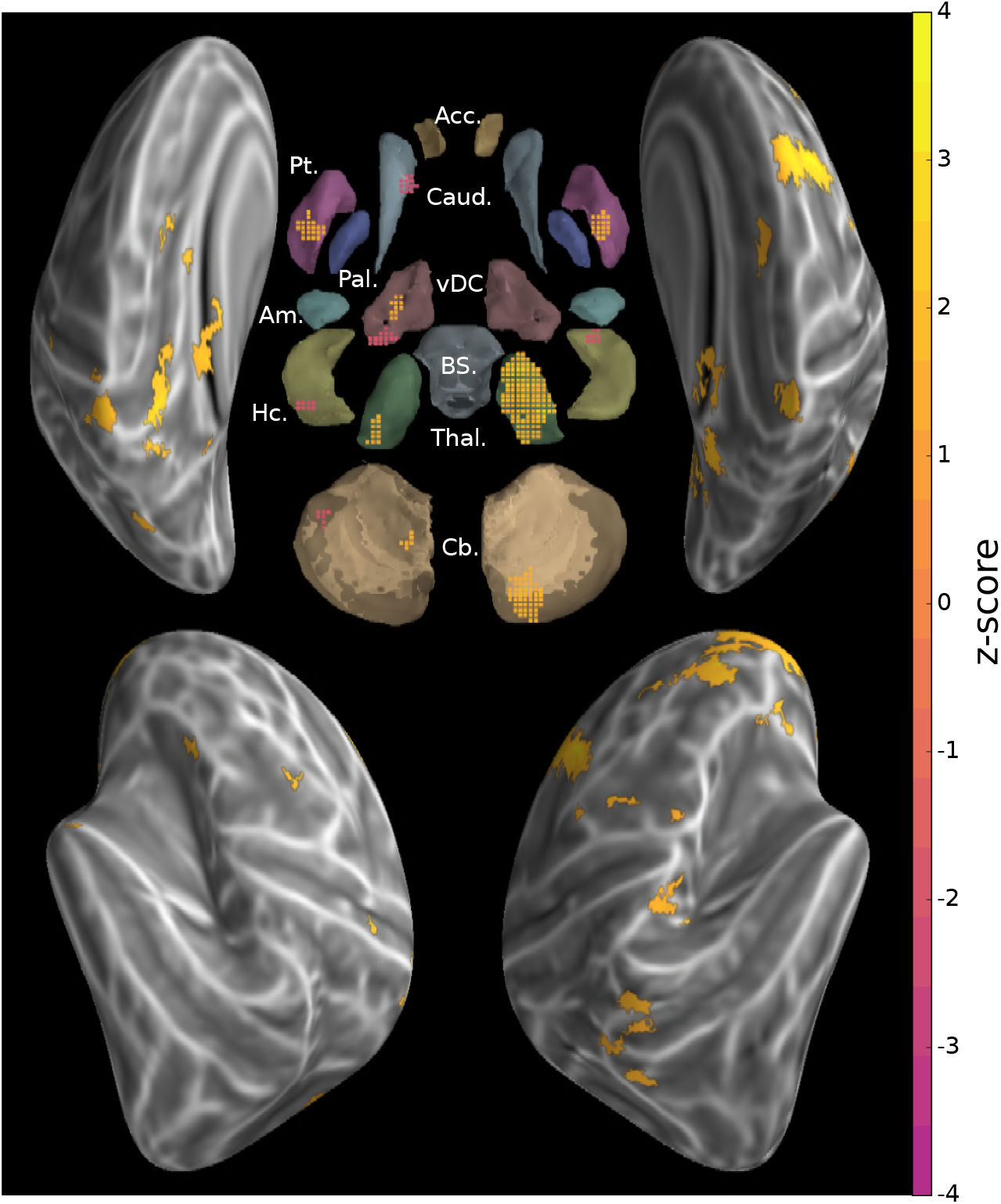
Conjunction of group searchlight contrast (paired t-test) between consolidated and new sequences discriminability maps and separate group discriminability maps for new and consolidated sequences (z-score thresholded at *p* < .05 TFCE-cluster-corrected) showing a reorganization of the distributed memory trace between these two stages; Acc.: Accumbens; Pt.:Putamen; Caud.:Caudate; Pal.:Pallidum; vDC:ventral Diencephalon; Am.:Amygdala; Hc.:Hippocampus; Thal.:Thalamus; Cb.:Cerebellum; BS:brain-stem

Discriminability between the consolidated sequences was significantly higher than that between the new sequences in bilateral sensorimotor putamen, thalamus and anterior insula, as well as in the ispilateral cerebellar lobule IX, posterior cingulate and parietal cortices, and contralaterally in the lateral and dorsal premotor, supplementary motor, frontopolar and dorsolateral prefrontal cortices in addition to cerebellar Crus I. By contrast, the pattern dissimilarity was higher for the new sequences in bilateral hippocampi as well as the body of the caudate nuclei, subthalamic nuclei, and cerebellar Crus II ipsilaterally. Although striatal activity patterns differentiating newly acquired sequences were found in contralateral putamen (fig. S1), this discriminability was significantly larger for consolidated sequences in sensorimotor regions of the putamen bilaterally.

## Discussion

In the present study, we aimed to identify the brain networks whose activity patterns differentiate between representations of multiple motor sequences during their execution in different phases of learning (newly learned vs consolidated). Using an MVPA approach, we considered that stable local patterns of activity could be used as proxy for the specialization of neuronal circuits supportive of the efficient retrieval and expression of sequential motor memory traces. To investigate the differential pattern strength, we computed novel unbiased multivariate distance and applied robust permutation-based statistics with adaptive cluster correction.

### A distributed network for the representation of finger motor sequence

Our results provide evidence for an extended network of brain regions that shows reliable discrimination of sequence-specific activity patterns for both the consolidated and novel sequences. At the cortical level, we found a network encompassing the supplementary motor and premotor areas as well as posterior parietal cortices bilaterally and contralateral somatosensory motor cortex. These findings are consistent with earlier MVPA investigations (Wiestler and Diedrichsen 2013; Nambu et al. 2015). Indeed, similar discriminative power of motor sequence representations within the ipsilateral premotor and parietal cortices has previously been described (Wiestler and Diedrichsen 2013;Waters-Metenier et al. 2014;Waters et al. 2017), notably when the non-dominant hand is used for fine dexterous manual skills. Interestingly, we also found significant neural representations for both learning stages in the contralateral primary motor and somatosensory (M1/S1) cortices, more specifically around the hand knob area (Yousry et al. 1997) for which finger somatotopy is measurable using fMRI (Ejaz et al. 2015). The latter results suggest that these primary cortical regions play a critical role in building experience-related motor sequence memory traces. Yet such an interpretation must be taken with caution, as it has recently been reported that the capacity to discriminate between sequences based upon signals from these regions could simply be due to the stronger activity evoked by the first finger press in the sequence, and not to activity from the whole finger sequence (Yokoi et al. 2017). Yet although conjectural, we do not believe that such an effect can explain our pattern of results because, while the newly learned sequences began with different fingers, both consolidated sequences were discriminated despite the fact that the first finger presses were the same. Finally, while being located around the hand knob, the spatial extent of the M1/S1 representation in our study was smaller compared to that found by Wiestler and Diedrichsen (2013). This may be due, however, to differences in our design, notably in the uninterrupted repetition of the motor sequence during practice, and in the fact that none of our sequences engaged the thumb, which has a more distinctive M1/S1 cortical representation than the individual fingers (Ejaz et al. 2015).

The conjunction of new and consolidated sequences discriminability maps further revealed that a common cortical processing network, including non-motor primary and associative regions, carries sequential information across learning stages, that can originate from visually presented instruction and short-term-memory to motor sequence production. Herein, the visual occipital cortices, likely reflecting processing of the visual stimuli as low-level visual mapping of shapes (Miyawaki et al. 2008;Pilgramm et al. 2016), as well as the ventro-temporal regions, known to support higher level Arabic number representation (Shum et al. 2013;Peters et al. 2015) were found to discriminate between sequences in both stages of learning (fig. 2). The dorsolateral prefrontal cortex (DLPFC), which also exhibited pattern discriminability, was suggested previously to process the sequence spatial information in working memory, preceding motor command (Robertson et al. 2001). In fact, we believe that the cognitive processing required in our task, implying notably to switch between sequences, to maintain them in working memory and to inhibit competing ones, could have magnified this frontal associative representation in our study.

In sum, the regions found to carry sequence information regardless of the learning phase in the present study show some overlap with the network known to be implicated in MSL, such as primary and secondary motor cortices, as typically revealed in activation-based studies (Doyon et al. 2009b; Dayan and Cohen 2011; Hardwick et al. 2013). However, we also found significant representations in the occipital, temporal and insular cortices. This discrepancy can be attributable to the shift from an activation-based inference to one based on the presence of sequential information in activity patterns, but also by the recruitment of additional regions for the processing of this information in stimuli and its maintenance in working memory required by the task.

### Cortico-subcortical representational reorganization underlying memory consolidation following MSL

By contrasting the maps of multivariate distances for consolidated and newly acquired sequences, we identified the networks that reveal increased versus decreased discriminability of sequential representations in the early stages of the MSL consolidation (fig. 3).

At the cortical level, we found that the contralateral premotor and bilateral parietal regions showed a stronger representation for consolidated sequences. This pattern likely reflects that the tuning of these neural populations to coordinated movements is consolidated early after learning (Pilgramm et al. 2016; Makino et al. 2017; Yokoi et al. 2017), as was previously observed when contrasting sequence that underwent a longer training to new ones (Wiestler and Diedrichsen 2013). Importantly, no significant changes in representational magnitude were found in the contralateral primary somatosensory cortex after consolidation. This is in line with the fact that M1 representational geometry has been shown to be strongly shaped by ecological finger co-activations (Ejaz et al. 2015), and to be resistant to extensive training of a sequence built on a new co-activation structure (Beukema et al. 2018). While the role of the motor cortex in MSL is undeniable, its plasticity in consolidation is still debated (Omrani et al. 2017). In fact, recent results revealed that after a M1 insult or even rapidly after M1 inactivation, a trained motor skill can still be expressed (Kawai et al. 2015; Bollu et al. 2018) arguing for its complementary, redundant and partially independent representation in subcortical regions.

Interestingly, significant differences at the subcortical level were found in bilateral putamen and more specifically in their sensorimotor regions. This is consistent with findings from activation studies that reported increased functional activity after consolidation in this structure (Debas et al. 2010, 2014; Albouy et al. 2013b; Fogel et al. 2017; Vahdat et al. 2017). Significant representational changes were also found in the bilateral thalami, and could reflect the relay of information between the cortex and cerebellum, striatum or spinal regions (Doyon et al. 2009a; Haber and Calzavara 2009). Finally, representation changes were detected in the cerebellum, including ipsilateral Lobule IX, shown to correlate with sequential skill performance (Orban et al. 2010;Tomassini et al. 2011) as well as contralateral Crus II which connectivity with prefrontal cortex is thought to support motor functions (Ramnani 2006). However, no significant difference was observed in Lobule V of the cerebellum that is known to carry finger somatotopic representations (Wiestler et al. 2011) and to show global activation during practice (Doyon et al. 2002).

Concurrently with the representational increase in the above-mentioned network, we found only a few disparate regions that showed decreased sequence discrimination, namely the caudate nuclei, subthalamic nuclei and cerebellar Crus II ipsilaterally as well as bilateral hippocampi. Hippocampal activation in early learning has formerly been hypothesized to support the temporary storage of novel explicitly acquired motor sequence knowledge and to contribute to the reactivations of the distributed network during offline periods and sleep in particular. Yet such contribution of the hippocampus has been shown to be progressively disengaging afterward (Albouy et al. 2013b), and thus our results are consistent with the idea of the hippocampus playing a transient supportive role in early MSL, notably in encoding sequential information (Davachi and DuBrow 2015). Our findings of a differential implication of dorsomedial and dorsolateral striatum in sequence representation during learning and expression of a mastered skill specifies the changes in activity in these regions in the course of MSL described by earlier studies (Lehéricy et al. 2005;François-Brosseau et al. 2009;Jankowski et al. 2009;Reithler et al. 2010; Corbit et al. 2017; Fogel et al. 2017; Kupferschmidt et al. 2017). Indeed, our results uncover that this shift in activity purports a genuine reorganization of circuits processing sequence-specific information, similar to what was reported at the neuronal level in animals (Miyachi et al. 2002;Costa et al. 2004;Yin et al. 2009).

While our results show that the topology of the network representing motor sequential information differs between consolidated and newly acquired memory traces, the present study was not designed to investigate the information-content of hippocampal, striatal or cerebellar sequence representations. These were previously assessed at cortical level for finger sequences (Kornysheva and Diedrichsen 2014;Wiestler et al. 2014) as well as for larger forearm movements (Haar et al. 2017). However, the hypothesized extrinsic and intrinsic skill encoding in the respective hippocampal and striatal systems (Albouy et al. 2013a) remains to be assessed with a dedicated experimental design similar to that used by Wiestler et al. (2014) to investigate such representations at the cortical level.

Importantly, our study investigated the change in neural substrates of sequence representation after limited training and following sleep-dependent consolidation. This is in contrast to previous investigations that studied sequences trained intensively for multiple days (Nambu et al. 2015) and compared their discriminability to that of newly acquired ones (Wiestler and Diedrichsen 2013). Therefore, in our study, the engagement of these representations for expressing the sequential skill may further evolve, strengthen or decline locally with either additional training or offline memory reprocessing supported in part by sleep.

### Methodological considerations

To limit the level of difficulty and the duration of the task, only four sequences were performed by participants, two consolidated and two newly acquired. This low number of sequence per condition could be a factor limiting the power of our analysis, as only a single multivariate distance is assessed for each of these conditions. Moreover, initial training sessions of the consolidated sequences were each comprised of a single sequence performed in blocks longer than in the present task, designed for multivariate investigation. The current task, by requiring additional cognitive resources (such as instruction processing, retention in working memory, switching and inhibition of other sequences), could have triggered some novel learning for the consolidated sequences. This seems unlikely however, as this was not reflected in performance changes throughout the task. The switching component could partly explain the pattern of results found here, as shifting between overlapping sets of motor commands has been shown to further implicate the dorsal striatum in collaboration with the prefrontal cortex (Monchi et al. 2006).

Another potential limitation relates to the fact that the present representational analysis disregarded the behavioral performance. Nevertheless, the chained non-linear relations between behavior, neural activity and BOLD signal were recently established to have limited influence on the representational geometry extracted from Mahalanobis cross-validated distance in primary cortex, sampled across a wide range of speed of repeated finger-presses and visual stimulation (Arbuckle et al. 2018). Therefore, despite behavioral variability and potential ongoing evolution of the memory trace, we assumed that the previously encoded motor sequence engrams were nevertheless retrieved during this task as supported by the significant differences in activity pattern discriminability and the persistent behavioral advantage observed for the consolidated sequences.

Finally, our results also entail that it is possible to investigate learning-related representational changes in a shorter time-frame and with less extended training than what was investigated before (Wiestler and Diedrichsen 2013; Nambu et al. 2015), including in subcortical regions where neuronal organization differs from that of the cortex. The use of a novel multivariate distance could have contributed to obtain these results by achieving increased sensitivity and statistical robustness (Walther et al. 2016).

## Conclusion

Our study shows that the consolidation of sequential motor knowledge is supported by the reorganization of newly acquired representations within a distributed cerebral network. We uncover that following learning, local activity patterns tuned to represent sequential knowledge are enhanced not only in extended cortical areas, similarly to those shown after longer training (Wiestler and Diedrichsen 2013), but also in dorsolateral striatum, thalamus and cerebellar regions. Conversely, a smaller network showed a decrease of sequence specific patterned activation after consolidation, occurring specifically in dorsomedial striatum that supports cognitive processing during early-learning (Doyon et al. 2018) as well as in the hippocampus which carries explicit encoding of motor sequential extrinsic representation (Albouy et al. 2013b;King et al. 2017) and play a significant role in the offline reprocessing. Despite discrepancies with GLM-based activity changes observed previously, the results of our novel representational approach corroborate their interpretations that the differential plasticity changes in the latter regions subtend MSL consolidation (Albouy et al. 2015). Importantly, these results reveal for the first time in humans that such changes are determined by the local implementation of distributed neural coding of sequential information. Yet such consolidation-related representational changes need to be further investigated through exploration of the dynamic mechanism mediating this sleep-dependent mnemonic process, which is known to reorganize progressively the cerebral network by repeatedly reactivating the memory trace (Fogel et al. 2017;Vahdat et al. 2017;Boutin et al. 2018).

## Materials and methods

### Participants

Right-handed young (*n* = 34,25 ± 6.2yr.) healthy individuals (19 females), recruited by advertising on academic and public website, participated in the study. Participants were excluded if they had a history of neurological psychological or psychiatric disorders, scored 4 and above on the short version of Beck Depression Scale (Beck et al. 1961), had a BMI greater than 27, smoked, had an extreme chronotype, were night-workers, had traveled across meridians during the three previous months, or were trained as musician or professional typist for more than a year. Their sleep quality was subjectively assessed, and individuals with score to the Pittsburgh Sleep Quality Index questionnaire (Buysse et al. 1989) greater or equal to 5, or daytime sleepiness Epworth Sleepiness Scale (Johns 1991) score greater than 9, were excluded.

Participants included in the study were also instructed to abstain from caffeine, alcohol and nicotine, to maintain a regular sleep schedule (bed-time 10PM-1AM, wake-time 7AM-10AM) and avoid taking daytime nap for the duration of the experiment. In a separate screening session, EEG activity was also recorded while participants slept at night in a mock MRI scanner and gradients sounds were played to both screen for potential sleep disorders and test their ability to sleep in the experimental environment;18 participants were excluded for not meeting the criterion of a minimum of 20min. in NREM2 sleep. After this last inclusion step, their sleep schedule was assessed by analyzing the data obtained from an actigraph (Actiwatch 2, Philips Respironics, Andover, MA, USA) worn on the wrist of the non-dominant hand for the week preceding as well as during the three days of experiment, hence certifying that all participants complied to the instructions.

Among the 34 participants, one did not show within-session improvement on the task, two didn’t sleep on the first experimental night, three were withdrawn for technical problems, one did not show up on first experimental session, one presented novel MRI contraindication. Thus, among the 26 participants that completed the research project, a group of 18 which, by design, followed the appropriate behavioral intervention for the present study, were retained for our analysis.

All participants provided written informed consent and received financial compensation for their participation. This study protocol was approved by the Research Ethics Board of the “Comité mixte d’éthique de la recherche - Regroupement en Neuroimagerie du Québec” (CMER-RNQ).

### Procedures and tasks

The present study was conducted over 3 consecutive evenings and is part of an experiment that aimed to investigate the neural substrates mediating the consolidation and reconsolidation of motor sequence memories during wakefulness and sleep that will be reported separately. On each day, participants performed the experimental tasks while their brain activity was recorded using MRI. Their non-dominant hand (left) was placed on an ergonomic MRI-compatible response pad equipped with 4-keys corresponding to each of the fingers excluding the thumb.

On the first day (D1), participants were trained to perform repeatedly a 5-element sequence (TSeq1: 1-4-2-3-1 where 1 indicate the little finger and 4 the index finger). The motor sequence was performed in blocks separated by rest periods to avoid fatigue. Apart for a green or a red cross displayed in the center of the screen, respectively instructing the participants to execute the sequence or to rest, there were no other visual stimuli presented during the task. Participants were instructed to execute the sequence repeatedly, and as fast and accurately as possible, as long as the cross was green. They were then instructed to rest for the period of 25 sec. as indicated by the red cross. During each of the 14 practice blocks, participants performed repeatedly 12 motor sequences (i.e. 60 keypresses per block). In case participants made a mistake during sequence production, they were instructed to stop their performance and to immediately start practicing again from the beginning of the sequence until the end of the block. After completion of the training phase, participants were then administered a short retention test about 15min later, which consisted of a single block comprising 12 repetitions of the sequence. Then the participants were scanned with concurrent EEG and fMRI for approximately two hours while instructed to sleep.

On the second day (D2), participants were first evaluated on the TSeq1 (1 block retest) to test their level of consolidation of the motor sequence, and were then trained on a new sequence (TSeq2: 1-3-2-4-1) which was again performed for 14 blocks of 12 sequences each, similarly to TSeq1 training on D1. Again, they were then scanned during sleep while EEG recordings were simultaneously acquired.

Finally, on the third day (D3), participants first performed TSeq1 for 7 blocks followed by 7 blocks of TSeq2, each block including 12 repetitions of the sequence or 60 keypresses. Following this last testing session, participants were then asked to complete an experimental task (here called MVPA task) specifically designed for the current study, similar to a previous study that investigated sequence representation by means of multivariate classification (Wiestler and Diedrichsen 2013). Specifically, participants performed short practice blocks of 4 different sequences, including TSeq1 and TSeq2 that were then consolidated, as well as two new finger sequences (NewSeq1: 1-2-4-3-1, NewSeq2: 4-1-3-2-4). In contrast to Wiestler and Diedrichsen (2013), however, all four sequences used only four fingers of the left-hand, excluding the thumb. Also, as for the initial training, sequences were instead repeated uninterruptedly and without feedback, in order to probe the processes underlying automatization of the skill.

Each block was composed of an instruction period of 4 seconds during which the sequences to be performed was displayed as a series of 5 numbers (e.g. 1-4-2-3-1), that could easily be remembered by the participant. The latter was then followed by an execution phase triggered by the appearance of a green cross. Participants performed 5 times the same sequence (or a maximum of 25 key-presses), before being instructed to stop and rest when the red cross was displayed.

The four sequences were assigned to blocks such as to include all possible successive pairs of the sequences using De-Bruijn cycles (Aguirre et al. 2011), thus preventing the systematic leakage of BOLD activity patterns between blocks in this rapid design. As a 2-length De-Bruijn cycle of the 4 sequences has to include each sequence 4 times, this yielded a total of 16 blocks. In our study, two different De-Bruijn cycles were each repeated twice in two separate scanning runs separated by approximately 5 minutes of rest, hence resulting in a total of 64 blocks (4 groups of 16 practice blocks for a total of 16 blocks per sequence). The blocks were synchronized to begin at a fixed time during the TR of the fMRI acquisition.

### Behavioral statistics

Using data from the MVPA-task, we entered the mean duration per block of correctly performed sequences into a linear mixed-effect model with a sequence learning stage (new/consolidated) by block (1-16) interaction to test for difference in their performance level, as well as the evolution during the task, with sequences and blocks as random effects and participants as the grouping factor. The same model was run with the number of correct sequences as the outcome variable. Two other models were also used on subsets of data to test separately if there was any significant difference in performance (speed and accuracy) between the two consolidated sequences and between the two new sequences. Full models outputs are reported in supplementary materials.

### MRI data acquisition

MRI data were acquired on a Siemens TIM Trio 3T scanner with two different setups. The first used a 32-channel coil to acquire high-resolution anatomical T1 weighted sagittal images using a Multi-Echo MPRAGE sequence (MEMPRAGE; voxel size=1mm isometric; TR=2530ms;TE=1.64,3.6,5.36,7.22ms;FA=7;GRAPPA=2;FoV=256 × 256 × 176*mm*) with the different echoes combined using a Root-Mean-Square (RMS).

Functional data were acquired with a 12-channel coil, which allowed to fit an EEG cap to monitor sleep after training, and using an EPI sequence providing complete cortical and cerebellum coverage (40 axial slices, acquire in ascending order, TR=2160ms;FoV=220 × 220 × 132mm, voxel size=3.44 × 3.44 × 3.3mm, TE=30ms, FA=90, GRAPPA=2). Following task fMRI data acquisition, four volumes were acquired using the same EPI sequence but with reversed phase encoding to enable retrospective correction of distortions induced by B0 field inhomogeneity.

### MRI data preprocessing

High-resolution anatomical T1 weighted images were preprocessed with Freesurfer (Dale et al. 1999; Fischl et al. 1999, 2008) to segment subcortical regions, reconstruct cortical surfaces and provide inter-individual alignment of cortical folding patterns. Pial and grey/white matter interface surfaces were downsampled to match the 32k sampling of Human Connectome Project (HCP) (Glasser et al. 2013). HCP subcortical atlas coordinates were warped onto individual T1 data using non-linear registration with the Ants software (Avants et al. 2008;Klein et al. 2009).

A custom pipeline was then used to preprocess fMRI data prior to analysis and relied on an integrated method (Pinsard et al. 2018) which combines slice-wise motion estimation and intensity correction followed by the extraction of BOLD timecourses in cortical and subcortical gray matter. This interpolation concurrently removed B0 inhomogeneity induced EPI distortion estimated by the FSL Topup tool using the fMRI data with reversed phase encoding (Andersson et al. 2003) acquired after the task. BOLD signal was further processed by detecting whole-brain intensity changes that corresponded to large motion, and each continuous period without such detected event was then separately detrended to remove linear signal drifts.

Importantly, the fMRI data preprocessing did not include smoothing, even though the interpolation inherent to any motion correction was based on averaging of values of neighboring voxels. This approach was intended to minimize the blurring of data in order to preserve fine-grained patterns of activity, with the resolution of relevant patterns being hypothetically at the columnar scale.

### Multivariate Pattern Analysis

#### Samples

Each block was modeled by two boxcars, corresponding to the instruction and execution phases respectively, convolved with the single-gamma Hemodynamic Response Function. Least-square separate (LS-S) regression of each event, which have been shown to provide improved activation patterns estimates for MVPA (Mumford et al. 2012), yielded instruction and execution phases beta maps for each block that were further used as MVPA samples.

#### Cross-validated multivariate distance

Similarly to Wiestler and Diedrichsen (2013) and Nambu et al. (2015), we aimed to uncover activity patterns that represented the different sequences performed by the participants. However, instead of calculating cross-validated classification accuracies, we opted for a representational approach by computing multivariate distance between activity patterns evoked by the execution of sequences, in order to avoid ceiling effect and baseline drift sensitivity (Walther et al. 2016). In the current study, we computed the cross-validated Mahalanobis distance (Nili et al. 2014;Diedrichsen et al. 2016;Walther et al. 2016), which is an unbiased metric that uses multivariate normalization by estimating the covariance from the GLM fitting residuals and regularizing it through Ledoit-Wolf optimal shrinkage (Ledoit and Wolf 2004). This distance, which measures discriminability of conditions, was estimated separately for pairs of sequences that were in a similar acquisition stage, that is, for the newly acquired and consolidated sequences.

#### Searchlight analysis

Searchlight (Kriegeskorte et al. 2006) is an exploratory technique that applies MVPA repeatedly on small spatial neighborhoods covering the whole brain while avoiding highdimensional limitation of multivariate algorithms. Searchlight was configured to select for each gray-matter coordinate their 64 closest neighbors as the subset of features for representational distance estimation. The neighborhood was limited to coordinates in the same structure (hemisphere or region of interest), and proximity was determined using respectively Euclidian and geodesic distance for subcortical and cortical coordinates. The extent of the searchlight was thus kept to such a local range to limit the inflation of false positive or negative results (Etzel et al. 2012, 2013).

#### Statistical testing

To assess statistical significance of multivariate distance and contrasts, group-level Monte-Carlo non-parametric statistical testing using 10000 permutations was conducted on searchlight distance maps with Threshold-Free-Cluster-Enhancement (TFCE) correction (Smith and Nichols 2009). The statistical significance level was set at *p* < .05 (with confidence interval ±.0044 for 10000 permutations) with a minimum cluster size of 10 features. TFCE enabled a locally adaptive statistics and cluster size correction that particularly fitted our BOLD sampling of sparse gray-matter coordinates, as well as the large differences in the sizes of the structures that were investigated.

The MVPA analysis was done using the PyMVPA software (Hanke et al. 2009) package with additional development of custom samples extraction, cross-validation scheme, efficient searchlight and multivariate measure computation, optimally adapted to the study design and the anatomy-constrained data sampling.

## Acknowledgments

We thank J. Diedrichsen for methodological advice on our multivariate representational analysis.

## Author contributions

1. Conceptualization: BP, AB, EG, HB, JD
2. Investigation: AB, EG, BP
3. Analysis: BP
4. Software development: BP
5. Writing: BP
6. Review and editing: BP, AB, EG, OL, HB, JD

## Supplementary materials

### Behavioral linear mixed-effect model outputs

Test for differences in speed as mean duration to perform a correct sequence per block

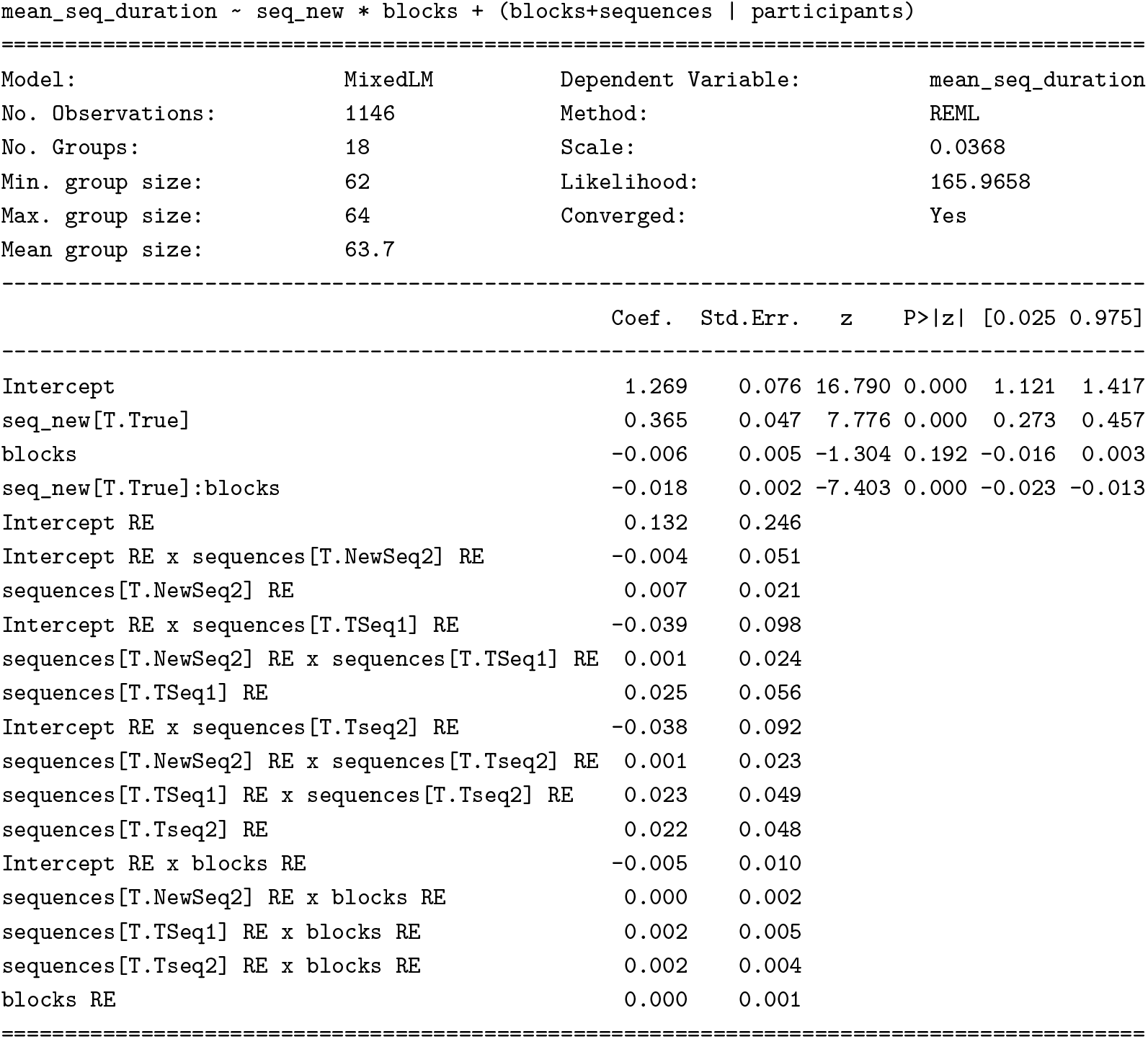

Test for differences in accuracy as the number of correct sequences over the 5 repetitions in a block

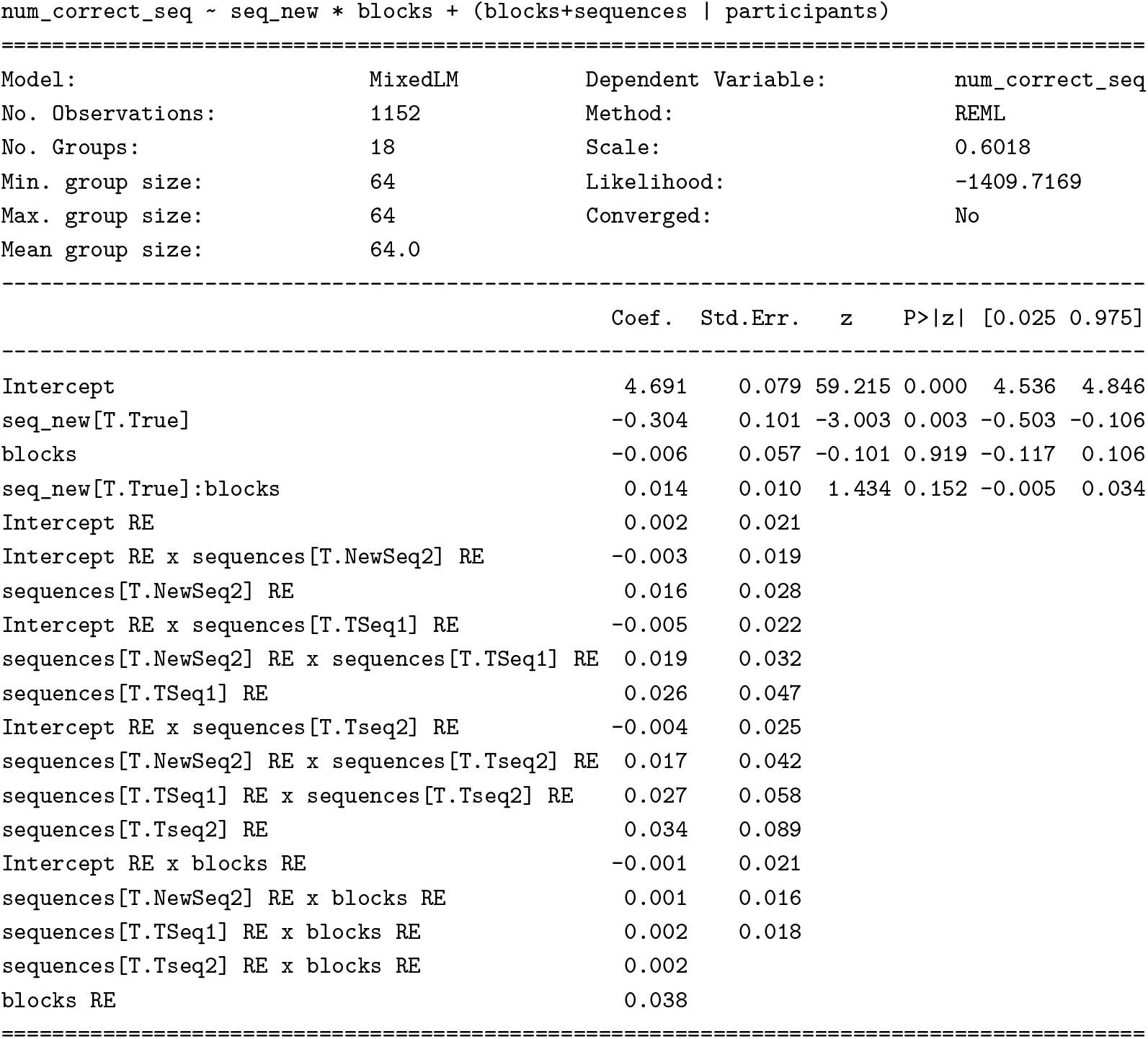

Test for differences in speed and accuracy between the new sequences

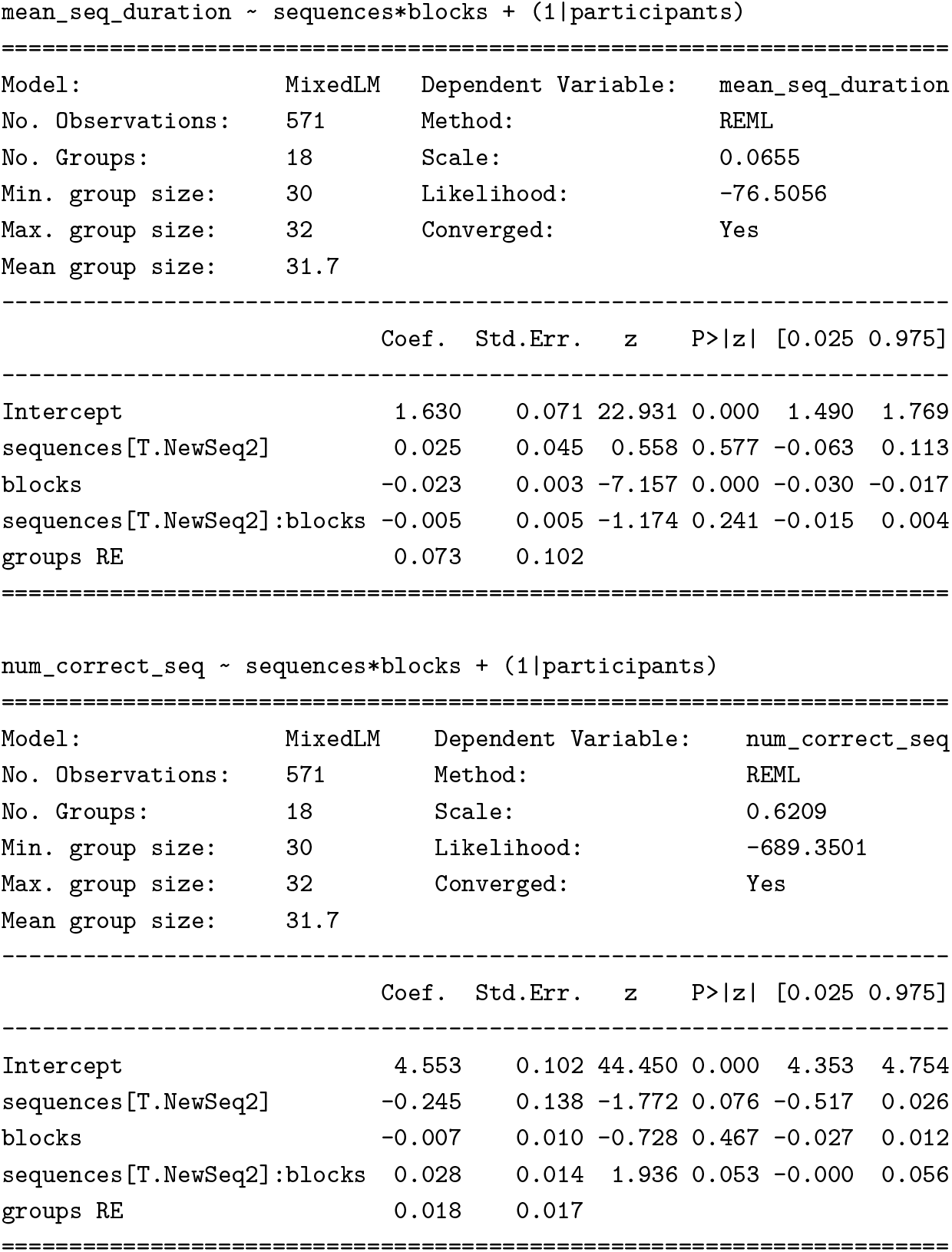

Test for differences in speed and accuracy between the consolidated sequences

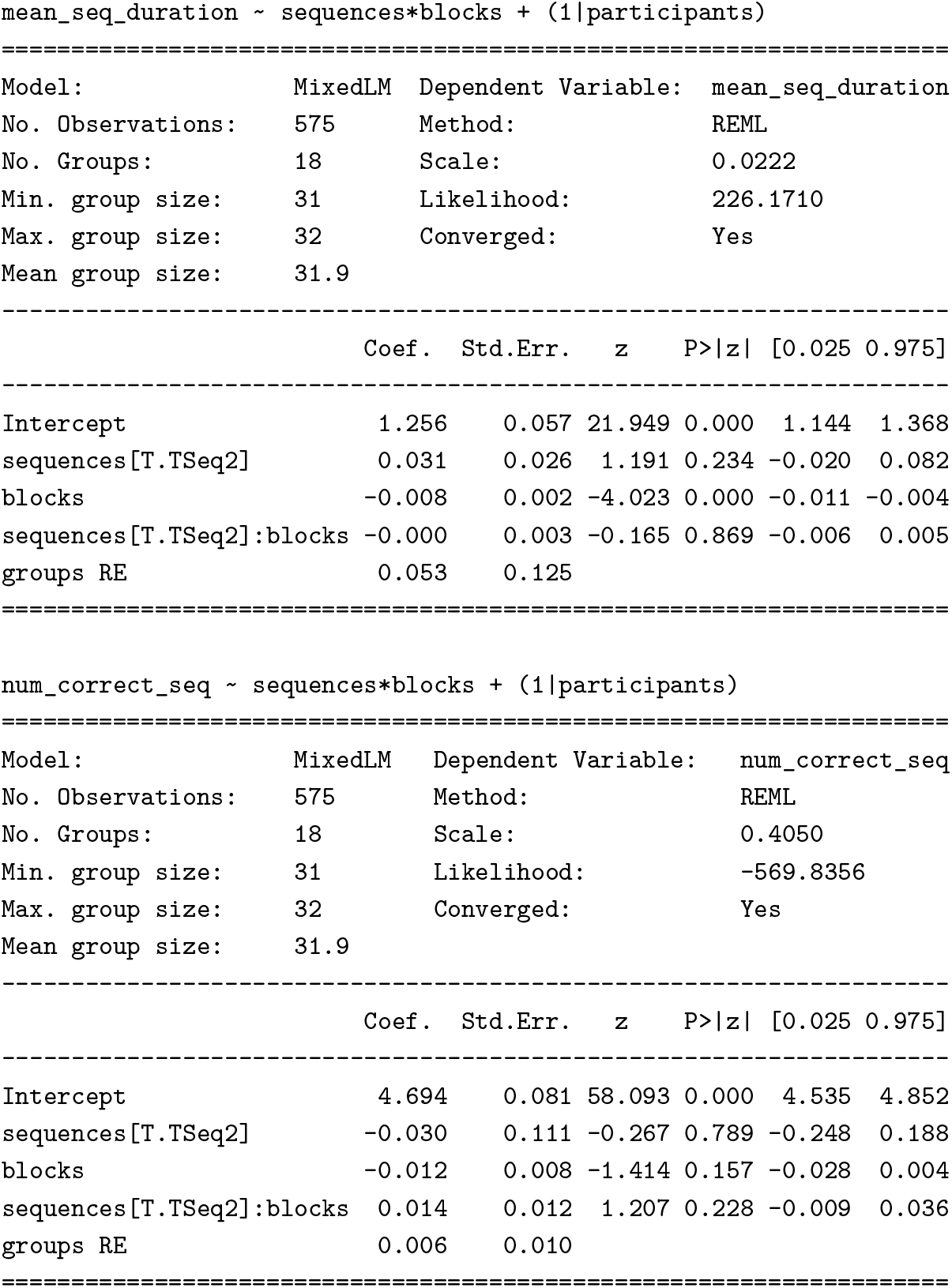

### Representational distance maps

**Figure S1.**
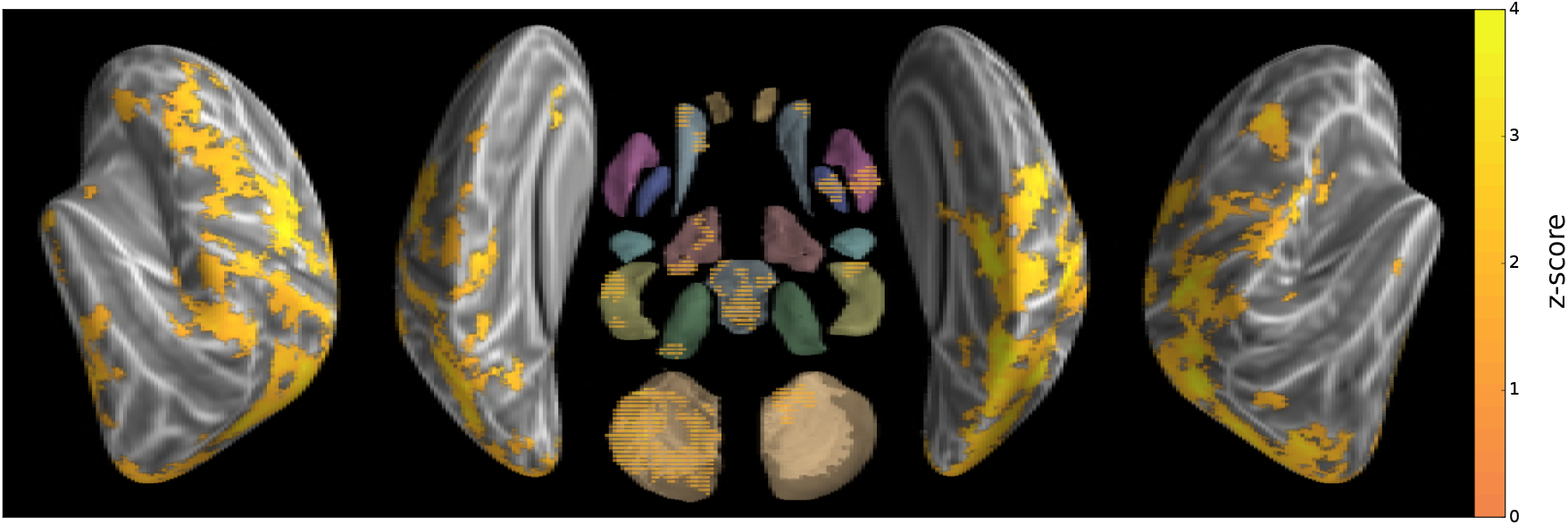
Group searchlight map of cross-validated Mahalanobis distance between the two new sequences (z-score thresholded at *p* < .05 TFCE-cluster-corrected)

**Figure S2.**
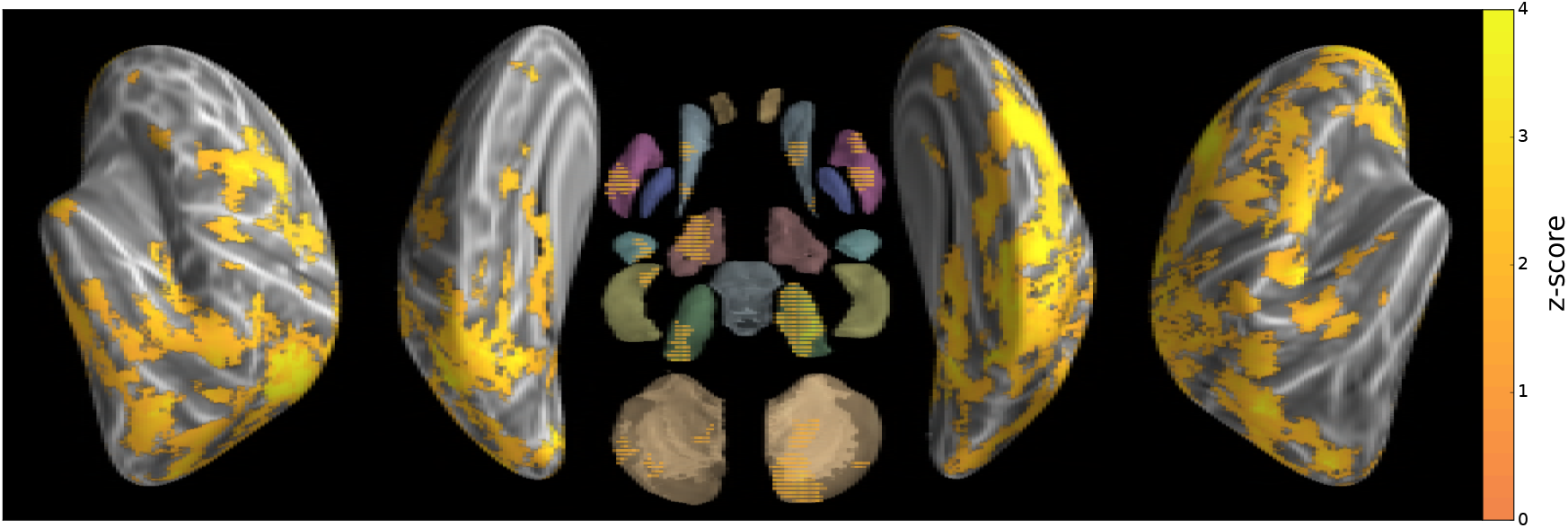
Group searchlight map of cross-validated Mahalanobis distance between the two consolidated sequences (z-score thresholded at *p* < .05 TFCE-cluster-corrected)

